# Discovery of synthetic lethal interactions from large-scale pan-cancer perturbation screens

**DOI:** 10.1101/810374

**Authors:** Sumana Srivatsa, Hesam Montazeri, Gaia Bianco, Mairene Coto-Llerena, Charlotte KY Ng, Salvatore Piscuoglio, Niko Beerenwinkel

**Affiliations:** Department of Biosystems Science and Engineering, ETH Zurich, 4058 Basel, Switzerland; SIB Swiss Institute of Bioinformatics, 4058 Basel, Switzerland; Department of Bioinformatics, Institute of Biochemistry and Biophysics, University of Tehran, Tehran, Iran; Institute of Pathology, University Hospital Basel, 4031 Basel, Switzerland; Department for BioMedical Research, University of Bern, 3008 Bern, Switzerland; Visceral Surgery Research Laboratory, Clarunis, Department of Biomedicine, University of Basel, Basel, Switzerland; Clarunis Universitäres Bauchzentrum Basel, Basel, Switzerland

**Keywords:** SLIdR, Synthetic lethality (SL), perturbation screen, driver genes, SL partner, causal inference, *AXIN1-URI1*

## Abstract

Despite the progress in precision oncology, development of cancer therapies is limited by the dearth of suitable drug targets^1^. Novel candidate drug targets can be identified based on the concept of synthetic lethality (SL), which refers to pairs of genes for which an aberration in either gene alone is non-lethal, but co-occurrence of the aberrations is lethal to the cell. We developed SLIdR (<u>S</u>ynthetic <u>L</u>ethal <u>Id</u>entification in <u>R</u>), a statistical framework for identifying SL pairs from large-scale perturbation screens. SLIdR successfully predicts SL pairs even with small sample sizes while minimizing the number of false positive targets. We applied SLIdR to Project DRIVE data^2^ and found both established and novel pan-cancer and cancer type-specific SL pairs. We identified and experimentally validated a novel SL interaction between *AXIN1* and *URI1* in hepatocellular carcinoma, thus corroborating the potential of SLIdR to identify new SL-based drug targets.

---

Key to exploiting SL in cancer therapy is the identification of a targetable dependent gene (SL partner) for a given genetically altered gene, such that a loss-of-function aberration in either gene alone does not affect cell viability, but aberrations in both genes are fatal to the cell (**Fig. 1a**). A classical example of SL in cancer therapy is the use of *PARP* inhibitors in *BRCA*-mutated cancers. The *BRCA1*/*2* genes involved in DNA double-strand break repair are often mutated in breast and ovarian cancers^3–5^, and hence such cancer cells rely on alternate DNA repair processes. *PARP1* plays a central role in these alternate DNA repair mechanisms^6,7^, and therefore inhibiting *PARP* results in catastrophic double-strand breaks during replication, ultimately leading to cancer cell death^8,9^.

**Fig. 1.**
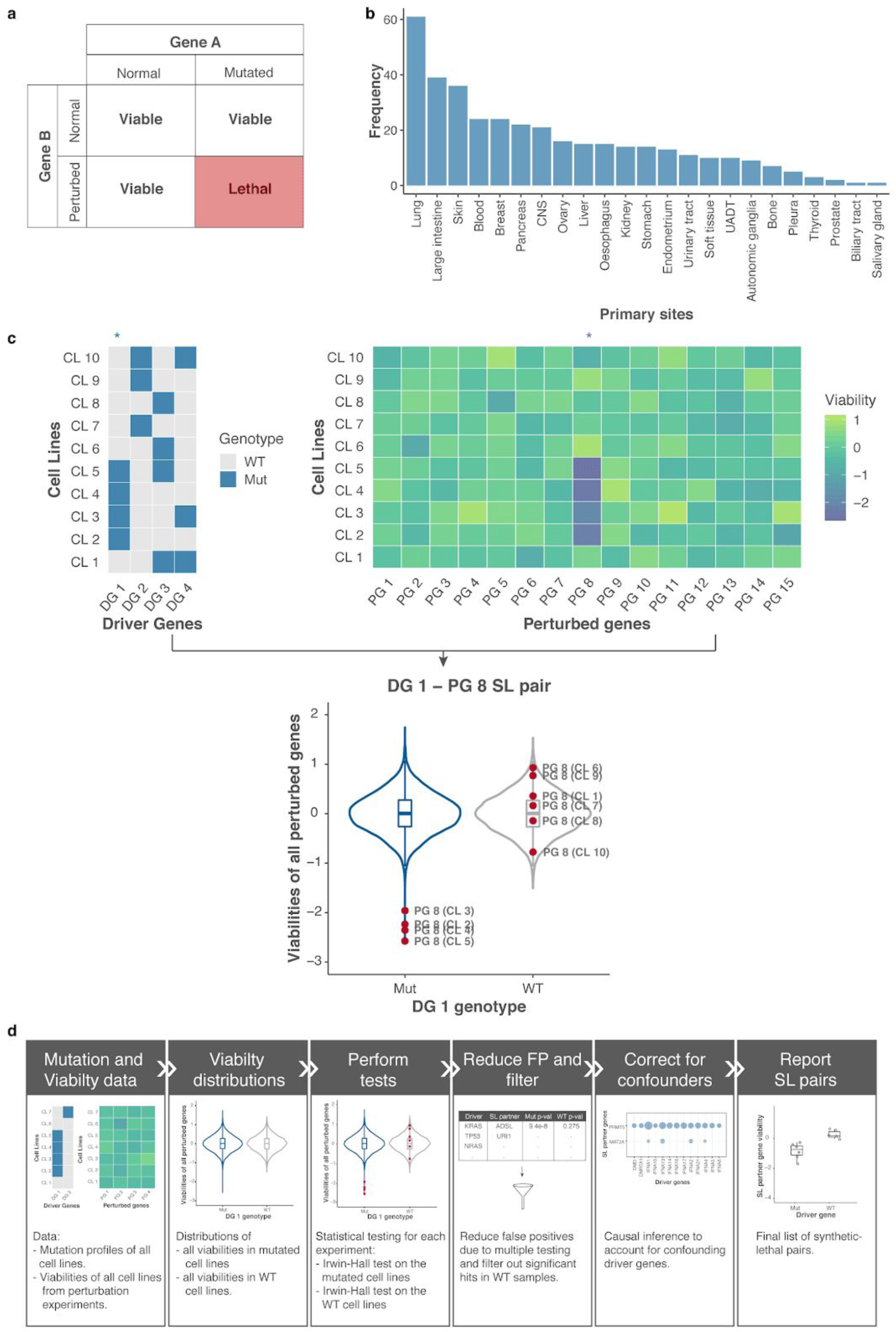
Overview and SLIdR workflow. **a**, Definition of a synthetic lethal pair: Aberration of gene A or knockdown of gene B alone do not affect the viability of the cell. However, the combination of mutated gene A and knockdown of gene B is lethal to the cell. **b**, Distribution of the number of cell lines with copy number data from CCLE across 23 different cancer types used in this study. **c**, Illustration of the SLIdR algorithm with a toy example. The data consists of driver genes DG 1-DG 4 and perturbed genes PG 1-PG 15 across cell lines CL 1-CL 10. Cell lines CL 2-5 are mutated in the driver gene DG 1 (Mut), while the remaining cell lines are DG 1 wild-type (WT). Comparison of viability distributions across all perturbed genes PG 1-PG 15 in the DG 1 mutated (Mut) and WT cell lines shows that perturbation of gene 8 (PG 8) results in reduced viability only in CL 2-5 and not the WT cell lines. Thus, PG 8 is a SL partner of DG 1. **d**, The computational pipeline illustrating the different steps performed to obtain the candidate SL pairs from mutation profiles and perturbation screen data.

In recent years, large-scale perturbation screens based on siRNA, shRNA, CRISPR, or small molecules in cell lines and organoids have been used to identify SL interactions. McDonald *et al.*^*2*^ conducted a large-scale deep RNAi screen targeting 7,837 genes in 398 Cancer Cell Line Encyclopedia^10^ (CCLE) models and provided a rich and robust dataset for the identification of SL pairs. However, the authors primarily analyzed gene interactions in a pan-cancer manner. Although pan-cancer analyses are statistically powerful due to their large sample sizes, the underlying data are diverse. We hypothesized that such rich large-scale perturbation screens can be exploited further to obtain SL pairs. We developed a novel method called SLIdR (<u>S</u>ynthetic <u>L</u>ethal <u>Id</u>entification in <u>R</u>) for predicting SL partners from such screens in both pan-cancer and cancer type-specific settings.

To define the set of genetically altered genes, we focused on significantly mutated genes reported by MutSig 2CV v3.1^11,12^ for each cancer type, and considered these genes to be altered in cell lines if they were subject to non-synonymous mutations or deep deletions (**Online methods**). We collectively refer to these altered genes as driver genes and these alterations as mutations. SLIdR aims to find SL partners for such drivers from perturbation data. We applied SLIdR to the Project DRIVE dataset^2^, focusing on cell lines from CCLE^10^ with available copy number data across various cancer types (**Fig. 1b**).

SLIdR adopts a rank-based statistical framework to robustly identify SL interactions between a driver gene and a perturbed gene (**Fig. 1c)**. In contrast to previous methods which perform statistical tests on the raw viability readouts^2,13^, SLIdR uses the normalized ranks of the viabilities across all perturbed genes for each cell line in order to increase statistical power for small sample sizes. For each driver gene, SLIdR first stratifies the cell lines into mutated and wild-type based on the mutation status of the driver gene. Subsequently, it tests, for each perturbed gene, whether the perturbation results in lower ranked viabilities in the mutated cell lines but not in the wild-type cell lines. SLIdR uses two Irwin-Hall tests to mine for such driver-perturbed SL gene pairs (**Online methods**). Cell lines with several co-occurring driver mutations can yield multiple SL pairs with the same perturbed gene. To identify the most likely SL pairs, we perform causal inference using matching-based potential outcome models. For a given candidate pair, we match the wild-type to mutated cell lines based on the other co-occurring mutations, thus achieving a covariate balance. Finally, SLIdR compares the viabilities of the matched groups and the significant SL pairs are reported (**Fig. 1d; Online methods**).

We first applied SLIdR to the DRIVE data in a pan-cancer setting. We identified 151 SL pairs (**Supplementary Table S1**) involving 84 driver genes (**Fig. 2a**). Out of the 151 SL pairs, five pairs involving *bona fide* driver genes *TP53, KRAS, BRAF, CTNNB1*, and *PIK3CA* exhibited self-dependency, i.e., paired with themselves as the SL partner gene. This proved to be an efficient quality check for our method as these are well-established drivers and their subsequent knockdown resulted in cellular mortality. We also found that some cell lines with several co-occurring mutations resulted in multiple driver genes pairing with the same SL partner (**Fig. 2b; Supplementary Fig. S1a**). For example, co-deletion of genes near p16 including *MTAP* and several interferons is common in several cancers and subsequently all these drivers paired with *MAT2A* as the SL partner. Using causal inference, we predicted the relevant driver genes for each SL partner (**Fig. 2c; Supplementary Fig. S1b**), resulting in 90 SL pairs across 42 driver genes (**Fig. 2d**).

**Fig. 2.**
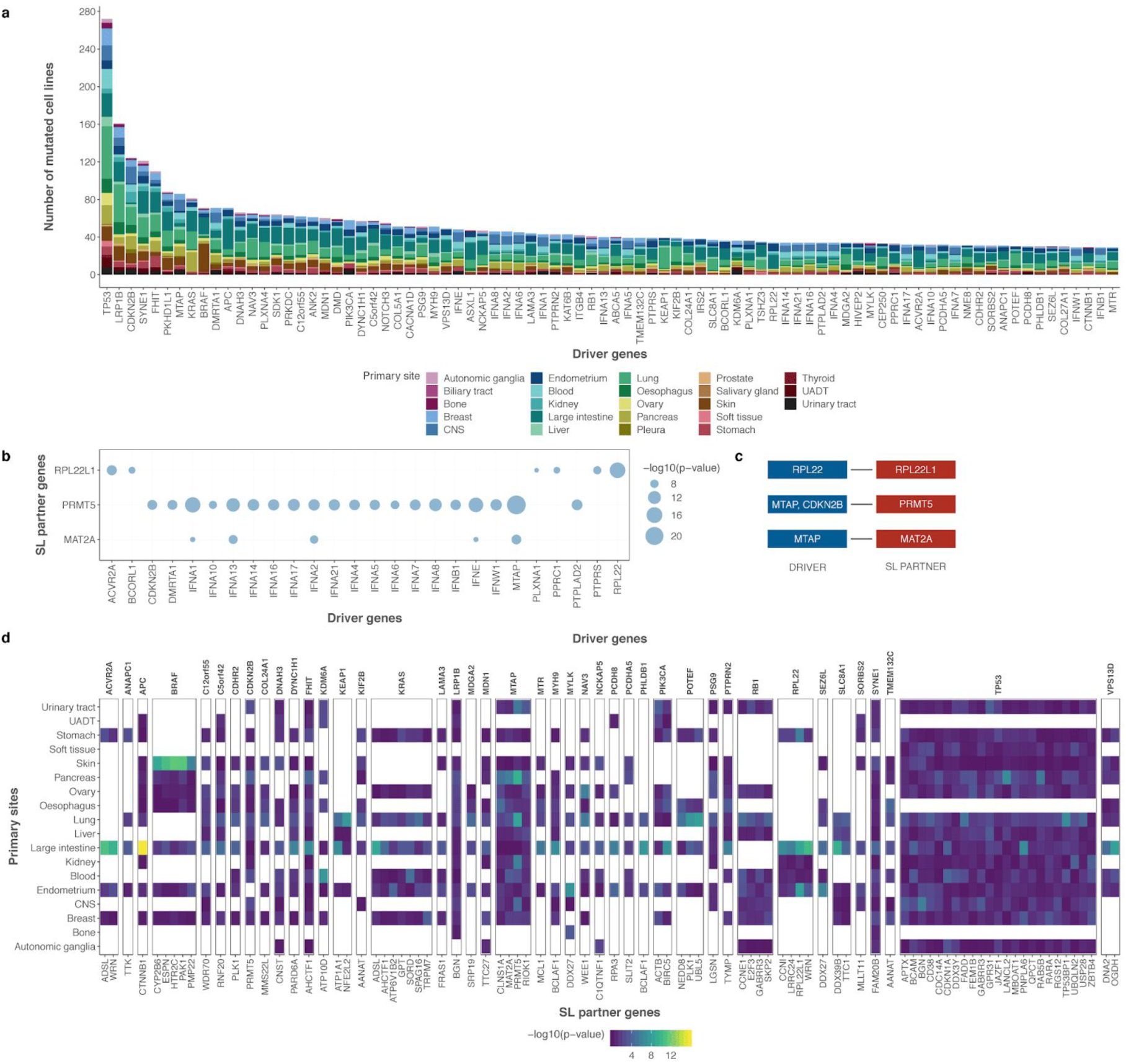
Pan-cancer SLIdR predictions. **a**, Stacked barplot indicating the frequencies of 84 mutated driver genes across different cancer types. **b**, Bubble-plot summarising the significance (-log10(p-value)) of different driver genes (x-axis) pairing with the same SL partner gene (y-axis). **c**, Corresponding list of significant SL pairs after accounting for confounding mutations and performing causal inference using matching-based potential outcome models. **d**, Differential sensitivities of pan-cancer SL pairs in subsets of cell lines grouped by primary sites (y-axis). Each panel corresponds to a specific driver gene (x-axis top) and encapsulates the significance profiles of all its SL-partners (x-axis bottom). Each column in a given panel depicts the significance profile of the SL pair in subsets of cell lines grouped by primary sites. The p-values are computed using IH-test.

Top predictions of SLIdR included *PRMT5, MAT2A*, and *RIOK1* as SL partners of *MTAP* which are all well-established vulnerable targets for *MTAP-*altered cells^2,14^. SLIdR also predicted *E2F3* and *SKP2* as SL partners for *RB1*^*2*^. Furthermore, *RPL22* showed lethality with its paralog *RPL22L1* confirming the findings of McDonald *et al.*^*2*^. *PIK3CA-BIRC5* was another reassuring pair as depletion of survivin (*BIRC5*) has been shown to have a pro-apoptotic effect in breast cancer cells with *PIK3CA* mutations^15,16^. In addition to established pairs, SLIdR also predicted several new SL pairs such as *KRAS-TRPM7* and *TP53-USP28*, which require further validation (**Supplementary Table S2**).

While pan-cancer analyses are favoured for their large sample sizes and the ability to identify shared targets across different cancer types, it is often difficult to identify such targets due to inherent differences between primary sites. The differential sensitivities of the predicted pan-cancer hits based on primary sites suggests that the majority of the signal is cancer type-specific (**Fig. 2d**). For example, SLIdR identified *NFE2L2* as the SL partner of the mutated *KEAP1*, both of which play an important role in cancer through Nrf2 pathway activation^17^. However, we noted that this SL interaction was largely due to signal from lung cancer samples (**Fig. 2d**), in accordance with Leiserson *et al.*^*18*^ who also reported the pair to be mutually exclusive in their pan-cancer TCGA analysis largely due to lung cancer samples. Thus, although a considerable number of pan-cancer hits are consistent with previous findings, this example among several others shows the need to identify cancer type-specific SL partners.

Consequently, we applied SLIdR to the DRIVE data in a cancer type-specific setting. For 17 cancer types, we identified a total of 839 SL pairs (**Supplementary Table S1**) over 233 unique driver genes. Out of the 233 drivers, 66 genes were mutated in more than one cancer type (**Fig. 3a**). However, the mutation profiles are diverse across cancer types, with *TP53* mutations being highly prevalent and observed in ∼81% of the cancer types, while well-known drivers such as *BRAF, APC*, and *PTEN* were distinctly associated with skin, large intestine, and endometrial cancers, respectively.

**Fig. 3.**
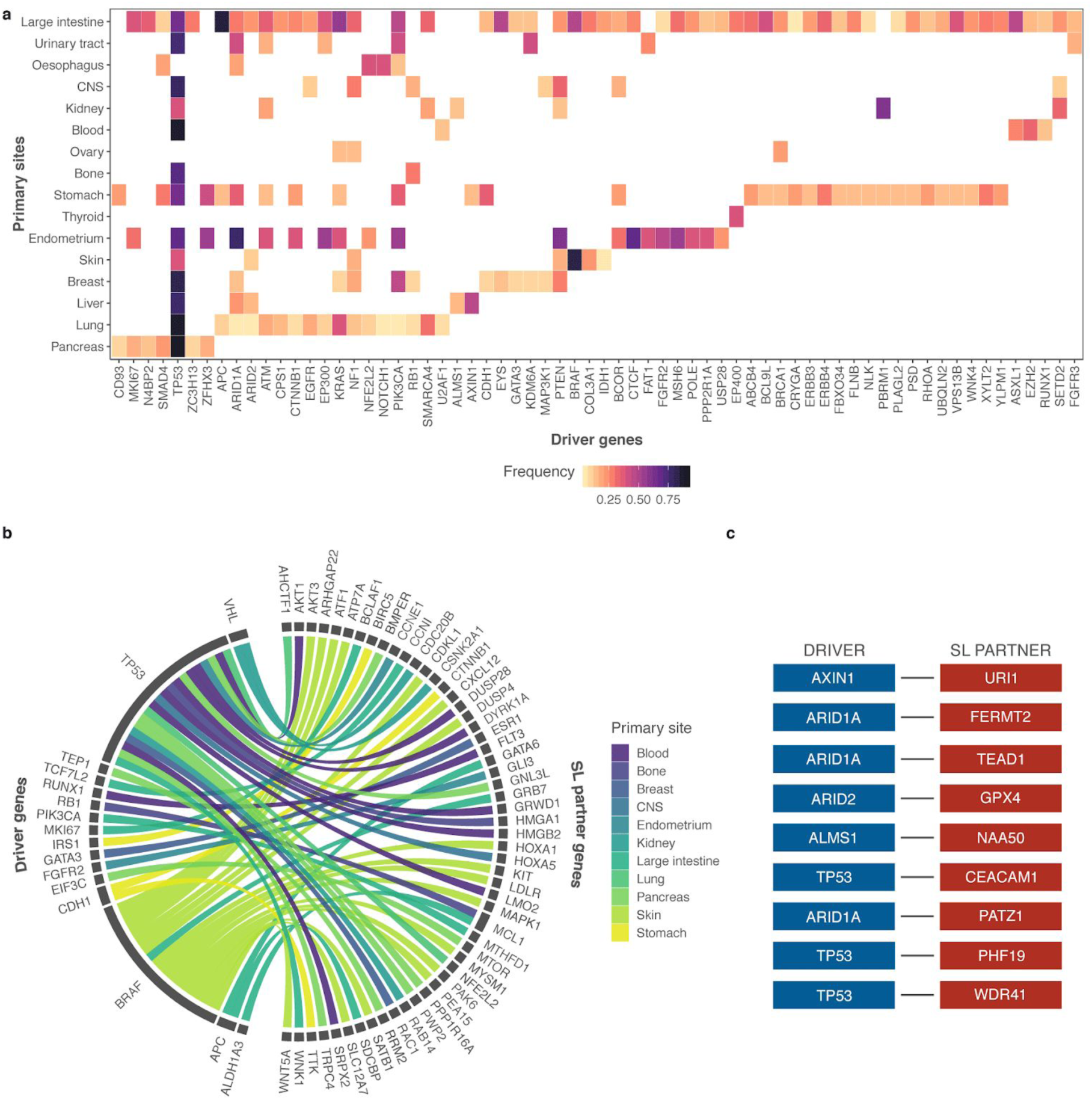
Cancer type-specific SLIdR predictions. **a**, Heatmap of frequencies of 66 driver genes across 16 cancer types. **b**, Circos plot summarizing the SL partners (right) of different driver genes (left) with literature evidence, across 11 cancer types. **c**, Top-ranked SL pairs in hepatocellular carcinoma reported by SLIdR.

Upon extensive literature survey of the SL pairs we identified 55 established and potential pairs with literature support (**Fig. 3b; Supplementary Table S2**). For example, SLIdR predicted *GATA3-ESR1* in breast cancer. *GATA3* is mutated in >10% of breast cancers and directly impacts *ESR1* enhancer accessibility, thereby altering binding potential and transcriptional targets in tumor cells^19^. Furthermore, *GATA3* mutations are almost never observed in ER-negative breast cancers, strongly suggesting SL. Despite the small sample size (7 cell lines), SLIdR also successfully elicited the *RB1-MCL1* pair in osteosarcomas. Loss-of-function *RB1* mutations are common in osteosarcomas^20,21^ and inhibition of *MCL1* has been shown to block tumor growth in osteosarcoma^22^. Additionally, SLIdR predicted several significant SL partners specific to *TP53*, including *TP53BP1, USP28, DDX3*, and *PNPLA6* in the pan-cancer setting, and *HMGA1, RAB14, and RAC1* in osteosarcoma, renal, and breast cancers, respectively (**Fig. 3b; Supplementary Table S2**). These examples highlight the ability of SLIdR to identify well-established and novel targets in both pan-cancer and cancer type-specific settings.

In hepatocellular carcinoma (HCC) we identified nine SL pairs (**Fig. 3c**). To demonstrate the predictive power of SLIdR, we sought to validate *AXIN1*-*URI1* (**Fig. 4a**), a novel pair and our top prediction in HCC. First, we validated the SL interaction between *AXIN1* and *URI1 in vitro* using SNU449, a HCC cell line carrying an *AXIN1* somatic mutation. Upon confirming that silencing of *URI1* using siRNA reduced *URI1* mRNA expression by >50% up to 96 hours post-transfection (**Fig. 4b**), we assessed cell viability by measuring cell proliferation rate. We observed that the knockdown of *URI1* in SNU449 cells significantly decreased proliferation compared to control cells (**Fig. 4c**).

**Fig. 4.**
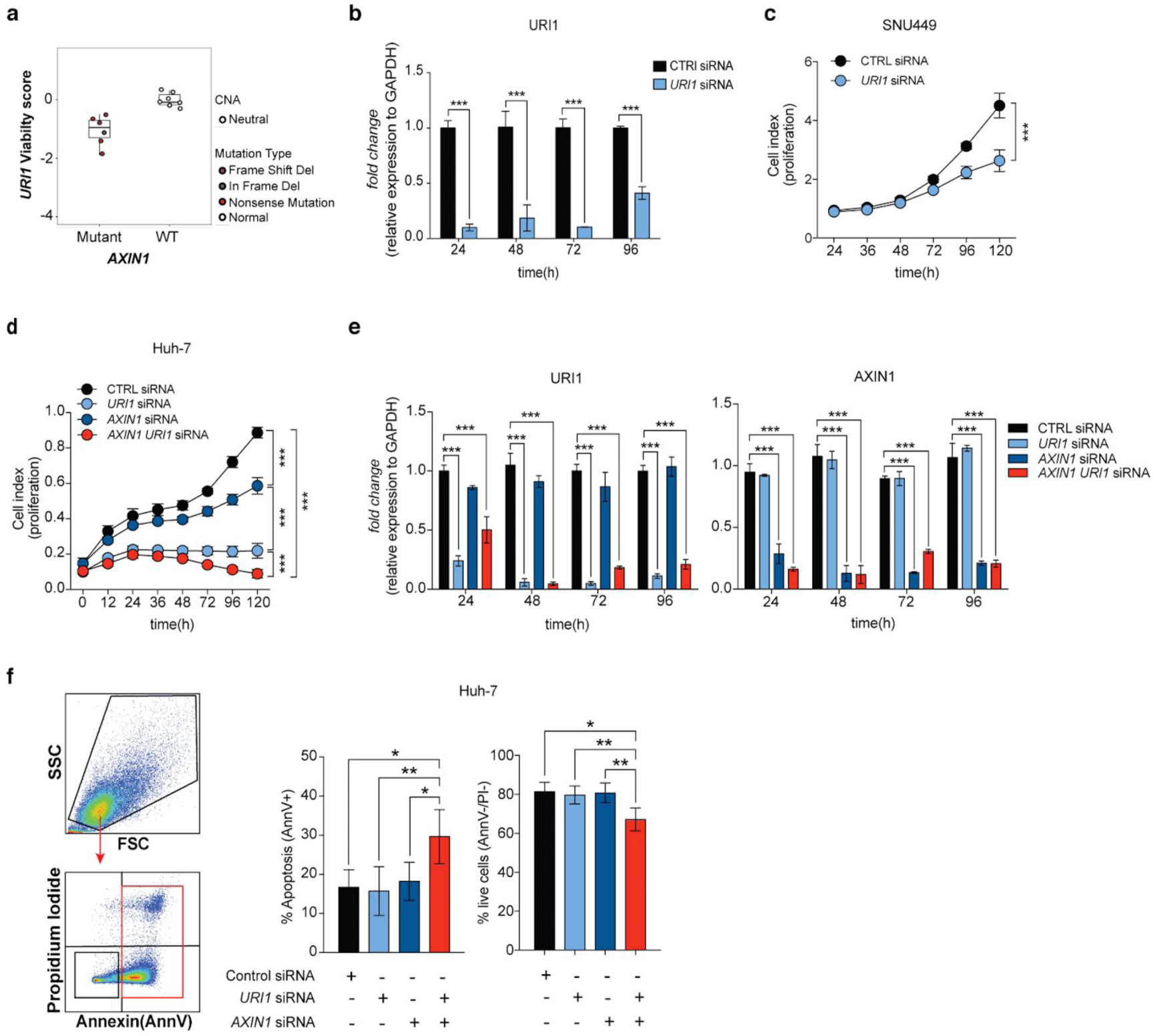
Functional validation of SL interaction between *AXIN1* and *URI1*. **a**, Top SL pair in HCC. Viability scores of *AXIN1* mutant vs wild-type (WT) HCC cell lines with *URI1* knockdown. **b**, RNA expression level (fold-change) of *URI1* relative to *GAPDH* in SNU449 cells transfected with control siRNA (black) or *URI1* siRNA (light blue). RNA levels were assessed by quantitative real-time PCR (qPCR). Error bars represent SD from two independent experiments. **c**, Cell proliferation assay in SNU449 cell line (*AXIN1* mutated) transfected with control siRNA (black) or *URI1* siRNA (light blue). Error bars represent standard deviation (SD) from two independent experiments. **d**, Cell proliferation assay in Huh-7 cell line (*AXIN1* WT) transfected with control siRNA (black), *URI1* siRNA (light blue), *AXIN1* siRNA (dark blue) or both (red). Error bars represent SD from three independent experiments. **e**, RNA expression levels (fold-change) of *URI1* (left) and *AXIN1* (right) relative to *GAPDH* in Huh-7 cell line transfected with control siRNA (black), *URI1* siRNA (light blue), *AXIN1* siRNA (dark blue) or both (red). Error bars represent SD from three independent experiments. **f**, Apoptosis assay using AnnexinV and propidium iodide (PI) staining in Huh-7 cell line (*AXIN1* wild-type) transfected with control siRNA (black), *URI1* siRNA (light blue), *AXIN1* siRNA (dark blue) or both (red). Quantification of the mean (+/-SD) percentage of apoptotic cells (AnnexinV+) and live cells (PI-/AnnexinV-) across the different groups (n=4) (right). Error bars represent SD from two independent experiments. For all experiments performed, statistical significance was assessed by multiple t-tests (* P < 0.05, ** P < 0.01, *** P < 0.001).

Next, we validated the SL interaction between *AXIN1* and *URI1* in Huh-7, an *AXIN1*-wildtype HCC-derived cell line. In this cell line, we silenced *AXIN1* and *URI1* alone or in combination. After confirming that silencing of the genes using siRNAs reduced their mRNA levels by 50-90% up to 96 hours post-transfection (**Fig. 4e**), we checked cells for growth inhibition using the same experimental approaches previously used in SNU449 (**Figs. 4d**,**e**). Huh-7 cells transfected with siRNAs targeting both *URI1* and *AXIN1* proliferated significantly less compared to cells transfected with CTRL siRNA, *AXIN1* siRNA or *URI1* siRNA alone (**Fig. 4d**). By staining cells with Annexin V and propidium iodide and analysing them by FACS, we showed that cells transfected with both *AXIN1* and *URI1* siRNA showed a higher proportion of apoptotic cells (15-20% more) and a lower proportion of living cells (20% less) compared to CTRL cells and cells transfected with either *URI1* or *AXIN1* siRNA alone (**Fig. 4f**), demonstrating that dual silencing was indeed fatal to the cells rather than merely arresting their proliferation.

To ensure that the cell death was truly due to synthetic lethality between *AXIN1* and *URI1* and not a result of an off-target effect or the cumulative cytotoxicity of the double siRNA transfection, we used the non-SL *AXIN1*-*TP53* pair as a negative control (**Supplementary Fig. S2a**). Dual silencing of *AXIN1* and *TP53* did not result in decreased cell proliferation compared to silencing of *TP53* or *AXIN1* alone (**Supplementary Fig. S2c**), confirming *AXIN1*-*URI1* as a novel SL pair in HCC.

Taken together, SLIdR provides a robust statistical framework to facilitate rapid discovery of SL interactions from large-scale perturbation screens in both pan-cancer and cancer type-specific settings. Particularly, in precision oncology, SLIdR can help in developing novel mutation-specific and effective personalised therapies.

## ONLINE METHODS

### Screening data

We used viability profiles published in Project DRIVE^2^, as well as corresponding mutation data and copy number data from the Cancer Cell Line Encyclopedia (CCLE) collection^10^ for 373 cell lines across 23 cancer types.

### Viability data from perturbation screens

Viability data specifies the viability of cell lines for each gene knockdown experiment. In Project DRIVE, 7837 genes were targeted by using an average of 20 pooled shRNAs per gene. The shRNA activities were defined as the quantile normalised log fold change in shRNA read counts 14 days after the start of the knockdown experiment to the shRNA abundance in the input library. The gene-level viability score of each cell line was computed by aggregation of shRNA activities using two computational methods, namely RSA^23^ and ATARiS^24^. The RSA method uses all shRNA reagents targeting a gene and can be used to identify essential, inert and active genes, while ATARiS only uses a subset of shRNAs with consistent activity across the cell lines and aims to provide a robust gene-level score by discarding shRNA reagents with off-target effects. ATARiS provides a relative score for the gene-level activity by median-centering the data for each reagent, and as a result, cannot distinguish between inert and essential genes.

To process the viability data, we removed essential genes using the RSA method as was performed in Project DRIVE^2^. Genes with an RSA value ≤-3 in more than 50% of cancer cell lines were reported as essential genes. In total, 460 and 185 genes were reported essential in cancer type-specific and pan-cancer settings, respectively. The resulting viability matrices consisted of the ATARiS scores for the remaining perturbed genes (rows) for each cell line (columns).

### Mutation and copy number data

For the pan-cancer setting, we focused on genes with mutations or copy-number aberrations in more than 30 cell lines. We downloaded mutation data and copy number data from the CCLE website, and binarized them as follows. A gene in a given cell line was assigned a value of 1 if it was subject to non-synonymous mutations, and 0 otherwise. For copy number data, we focused only on homozygous deletions and binarized a gene in a given cell line by assigning a value of 1 if the gene was homozygously deleted and a value of 0 otherwise. Finally, combining both these data, a driver gene in a given cell line was assigned a value of 1 if it was subject to non-synonymous mutations, deep deletions, or both; 0 otherwise.

In the cancer type-specific setting, to define the set of driver genes, we first used the MutSig 2CV v3.1^11,12^ MAF file from TCGA for each cancer type and focused only on significantly mutated genes (q <= 0.05). Next, we concentrated on genes with non-synonymous mutations in two or more cell lines, and excluded copy number data as it was very noisy in this setting. Thus, a gene in a given cell line was assigned a value of 1 if it was subject to non-synonymous mutations, and 0 otherwise. The resulting binarised mutation matrices described the mutation profiles for each cell line (column) across all driver genes (rows).

### SLIdR algorithm

SLIdR is a rank-based statistical framework to identify the presence of synthetic lethal dependency between a driver gene *d* and a perturbed gene *g*. For each driver gene *d*, we divided the cell lines into two groups according to the mutation status of *d*, namely wild-type cell lines (WT) and mutated cell lines (Mut). Further, we ranked the perturbed genes by their ATARiS scores, for each mutated and WT cell line and normalized it between 0 and 1. Due to a large number of perturbed genes, the normalized ranks have many distinct levels and are highly fine-grained. Hence, we assumed the normalized ranks to be continuous.

Let (*d, g*) be a fixed pair of driver and perturbed gene and *C*_*d*_ be the set of cell lines mutated in *d* of cardinality *n*. If (*d, g*) is an SL pair, based on the aforementioned definition (**Fig. 1a**), a mutation in driver gene *d* in combination with knockdown of gene *g*, results in low viabilities in mutated cell lines *C*_*d*_. We used a one-sided statistical test based on the Irwin-Hall distribution to test whether the viabilities of mutated cell lines *C*_*d*_ from knockdown of gene *g* are lower than expected by chance. We defined the null hypothesis *H*_0_ as the knockdown of gene *g* having no impact on the viability of the cell lines in *C*_*d*_. For each cell line *c* ∈ *C*_*d*_, we computed the normalized rank of the viability of *c* from knockdown of gene *g* across all other gene knockdowns in cell line *c*, and denoted this rank as *r*_*c*|*g*_. Under the null hypothesis, the normalized ranks take uniform random values in the interval [0, 1], *r*_*c*|*g*_ ∼ *U* (0, 1). The test statistic *T* for the pair (*d, g*) is then defined as the sum of normalized viability ranks of mutated cell lines *C*_*d*_ perturbed in gene *g*, 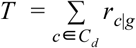. Under *H*_0_, the test statistic *T* is the sum of *n* independent uniform random variables on the unit interval and hence it follows the Irwin-Hall distribution of order *n*. The resulting p-value was computed as the lower tail probability *P* (*T* < *t*_*obs*_), where *t*_*obs*_ is the observed test statistic. For large *n*, computation of the Irwin-Hall probability distribution is either computationally expensive or numerically unstable. Therefore, we used the approximation *T* ∼ *N* (*n*/2, *n*/12) for *n* > 20.

Conversely, based on the definition of synthetic lethality (**Fig. 1a**), wild-type cell lines with respect to driver gene *d* are expected to behave similar to healthy cells when perturbed in gene *g*. Therefore, it is important to filter out genes which upon knockdown adversely alter the viabilities of WT cell lines. We used a two-sided Irwin-Hall test to filter out any pair (*d, g*) that reached statistical significance (α = 0.1) in the WT cell lines. However, we did not use this filter for pan-cancer setting due to the diverse nature of the cell types and cancer types.

### Multiple testing

We reduced the number of false positives arising from multiple testing by choosing a significance level of 1/(*M* × *N*), where *M* is the number of knockdowns and *N* is the number of driver genes. Therefore, we expect on average one false positive hit among all reported SL hits for each cancer. Our method is computationally inexpensive as it avoids performing all *M* × *N* tests. For each driver, we compute the test statistic for all perturbed genes and sort them in ascending order. The pre-ordering of the test statistics enables us to test for genes until the corresponding p-value is less than the chosen significance level. Further, we note that this approach was in good agreement with controlling the false discovery rate at 10% (**Supplementary Fig. S1c**).

### Causal inference

Cell lines are often subject to mutations or aberrations in multiple driver genes, and as a result, different driver genes pair with the same SL partner gene (**Fig. 2b; Supplementary Fig. S1a**). This is typically not an issue in the cancer type-specific setting but is prevalent in the pan-cancer setting. In order to identify the most likely SL pairs from the many confounding driver genes, we used matching-based potential outcome models. The main goal of the matching method is to emulate a randomized experiment by matching samples of treated and control groups according to covariates, thereby obtaining similar covariate distributions across the two groups. For a given driver gene *d*, the cell lines mutated in *d* constituted the treated group and the cell lines wild-type in *d* formed the control group. The remaining driver genes formed the confounding covariates and the viability of the SL partner gene *g* was used as the response or outcome variable. We used the *Matching* R package^25^ and performed propensity-score matching with a caliper of 0.1. Since matching is dependent on the order of the samples, we reshuffled and repeated matching 50 times. After each run, we recorded the standardised mean difference (smd) between the two groups for all covariates and chose the run with the lowest sum of smd across all covariates. Finally, for the chosen run, we performed a paired t-test between the responses of treated and control groups. We repeated this entire process for all the driver genes pairing with the same SL partner gene *g* and reported the significant (α = 0.05) SL pairs (**Fig. 2c; Supplementary Fig. S1b**).

### Code and data availability

The code base of SLIDR is available at https://github.com/cbg-ethz/slidr/. The raw shRNA data has already been published as a part of project DRIVE (https://data.mendeley.com/datasets/y3ds55n88r/4) and all the mutation and copy number data from CCLE is available at https://portals.broadinstitute.org/ccle. The MutSig 2CV v3.1^11,12^ MAF file for each cancer type is available at http://firebrowse.org/.

### Cell lines maintenance

Liver cancer-derived cell lines SNU449 and Huh-7 were obtained from the Laboratory of Experimental Carcinogenesis (Bethesda, MD, USA), authenticated by short tandem repeat profiling as previously described^26^ and tested for mycoplasma infection using a PCR-based test (ATCC). All cell lines were maintained under the conditions recommended by the provider. Briefly, all cell lines were cultured in DMEM supplemented with 5% Fetal Bovine Serum (FBS), non-essential amino-acids (NEAA) and antibiotics (Penicillin/Streptomycin). The cells were incubated at 37°C in a humidified atmosphere containing 5% CO_2_. Exponentially growing cells were used for all *in vitro* studies.

### Transient gene knockdown by siRNAs

Transient gene knockdown was conducted using ON-TARGET plus siRNA transfection. ON-TARGET plus SMARTpool siRNAs against human *URI1, AXIN1* and *TP53*, ON-TARGET plus SMARTpool non-targeting control and DharmaFECT reagent were all purchased from GE Dharmacon (**Supplementary Table S3**). Transfection was performed according to the manufacturer’s protocol. Briefly, log-phase liver cancer cells were seeded at approximately 60% confluence. Since residual serum affects the knockdown efficiency of ON-TARGET plus siRNAs, growth medium was removed as much as possible and replaced by serum-free medium (Opti-MEM; **Supplementary Table S3**). siRNAs were added to a final concentration of 25 nM. siRNAs targeting different genes can be multiplexed. Cells were incubated at 37°C in 5% CO_2_ for 24-48-72 hours (for mRNA analysis) or for 48-72 hours (for protein analysis). In order to avoid cytotoxicity, transfection medium was replaced with complete medium after 24 hours.

### RNA extraction and relative expression by qRT-PCR

Total RNA and proteins were extracted from cells at 75% confluence using TRIZOL (**Supplementary Table S3**) according to manufacturer’s guidelines. cDNA was synthesized from 1 μg of total RNA using SuperScript™ VILO™ cDNA Synthesis Kit (**Supplementary Table S3**). All reverse transcriptase reactions, including no-template controls, were run on an Applied Biosystem 7900HT thermocycler. Gene expression was assessed by using FastSart Universal SYBR Green Master Mix (**Supplementary Table S3**) and all qPCR were performed at 50°C for 2 min, 95°C for 10 min, and then 40 cycles of 95°C for 15 s and 60°C for 1 min on a QuantStudio 3 Real-Time PCR System (Applied Biosystems). The specificity of the reaction was verified by melting curve analysis. Measurements were normalized using GAPDH level as the reference. The fold change in gene expression was calculated using the standard ΔΔCt method as previously described^27^. All samples were analyzed in triplicates.

### Proliferation assay

Cell proliferation was assayed using the xCELLigence system (RTCA, ACEA Biosciences, San Diego, CA, USA). Background impedance of the xCELLigence system was measured for 12 s using 50 μl of room temperature cell culture media in each well of E-plate 16. Cells were grown and expanded in tissue culture flasks as previously described. After reaching 75% confluence, the cells were washed with PBS and detached from the flasks using a short treatment with trypsin/EDTA. 5000 cells were dispensed into each well of an E-plate 16. Growth and proliferation of the cells were monitored every 15 min up to 120 hours via the incorporated sensor electrode arrays of the xCELLigence system, using the RTCA-integrated software according to the manufacturer’s parameters. In the case of transient siRNA transfection, cells were detached and plated on xCELLigence 24 hours post-transfection.

### Apoptosis analysis by Flow Cytometry

Cells were collected 72 hours post siRNA transfection, stained with annexin V (FITC conjugate; **Supplementary Table S3**) and propidium iodide (PI), and analyzed by flow cytometry using the BD FACSCanto II cytometer (BD Biosciences, USA). Briefly, cells were harvested after incubation period and washed twice by centrifugation (1,200 g, 5 min) in cold phosphate-buffered saline (DPBS; **Supplementary Table S3**). After washing, cells were resuspended in 0.1 mL AnnV binding buffer 1X (ABB 5X, **Supplementary Table S3**) containing fluorochrome-conjugated AnnV and PI (PI to a final concentration of 1 ug/mL) and incubated in darkness at room temperature for 15 min. Following immediately, cells were analyzed by flow cytometry, measuring the fluorescence emission at 530 nm and >575 nm. Data were analyzed by FlowJo software version 10.5.3 (https://www.flowjo.com/).

For details on the reagents used, please refer to **Supplementary Table S3**.

## ACKNOWLEDGEMENTS & FUNDING

This work was partly supported by ERC Synergy Grant 609883 to N.B. Swiss Cancer League [KFS-3995-08-2016] and the Swiss National Foundation [PZ00P3_168165] to S.P.

## AUTHOR’S CONTRIBUTION

S.S. and N.B. conceived the presented idea. S.S. and H.M. developed and implemented the method and built the R package. N.B. supervised the computational study. S.P. and C.K.Y.N. supervised the experimental study. S.S. performed the downstream bioinformatics analyses and contributed to the visualizations. G.B. and M. C-L performed *in vitro* experiments. N.B., S.P. and C.K.Y.N. critically discussed the results. S.S., H.M., G.B., M. C-L, C.K.Y.N., S.P., and N.B. interpreted the results and wrote the manuscript. All authors agreed to the final version of the manuscript.

