## Supplementary Information for "Discovery of synthetic lethal interactions from large-scale pan-cancer perturbation screens"

**Supplementary tables**

**Supplementary Table S1:** Significant SL pairs identified by SLIdR in the pan-cancer and cancer type-specific analyses of DRIVE data.

**Supplementary Table S2:** Literature evidence for established and potential SL pairs predicted by SLIdR in pan-cancer and cancer type-specific analyses.

**Supplementary Table S3:** Reagents used in the functional validation of SL interaction between *AXIN1* and *URI1* and in the cross-validation of non-lethal interaction between *AXIN1* and *TP53* as negative control.

### Supplementary figures and figure legends

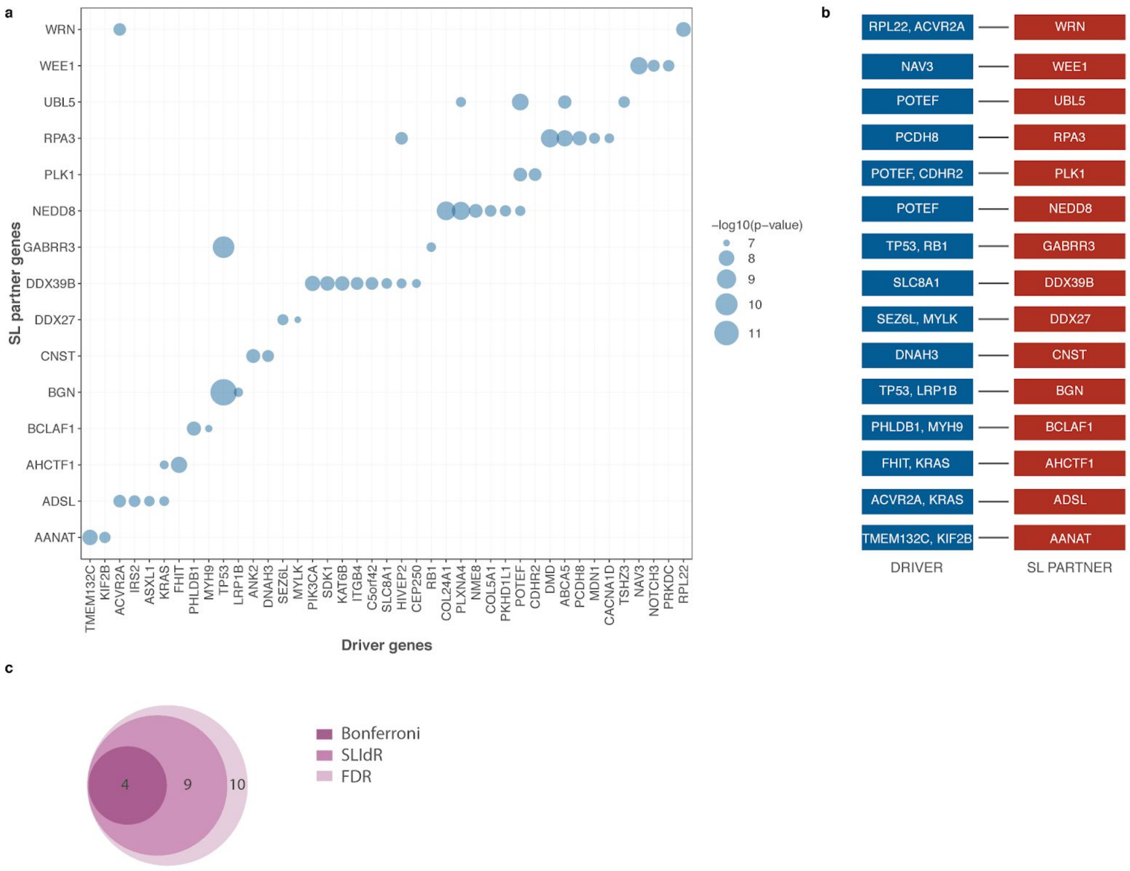

**Supplementary Fig. S1. Confounding in pan-cancer analysis and comparison of results from different multiple testing methods. a**, Bubble-plot summarising the significance (-log10(p-value)) of different pairs of driver genes (x-axis) and their SL partner genes (y-axis) in the pan-cancer analysis. **b**, Corresponding list of significant SL pairs after accounting for confounding mutations and performing causal inference using matching-based potential outcome models. **c**, An example illustrating the consensus between SLIdR hits and hits resulting from controlling the false discovery rate (FDR) at 10%. Venn-diagram comparing

SLIdR hits and hits reported after multiple testing corrections with FDR (10%) and Bonferroni (10%) methods in liver cancer.

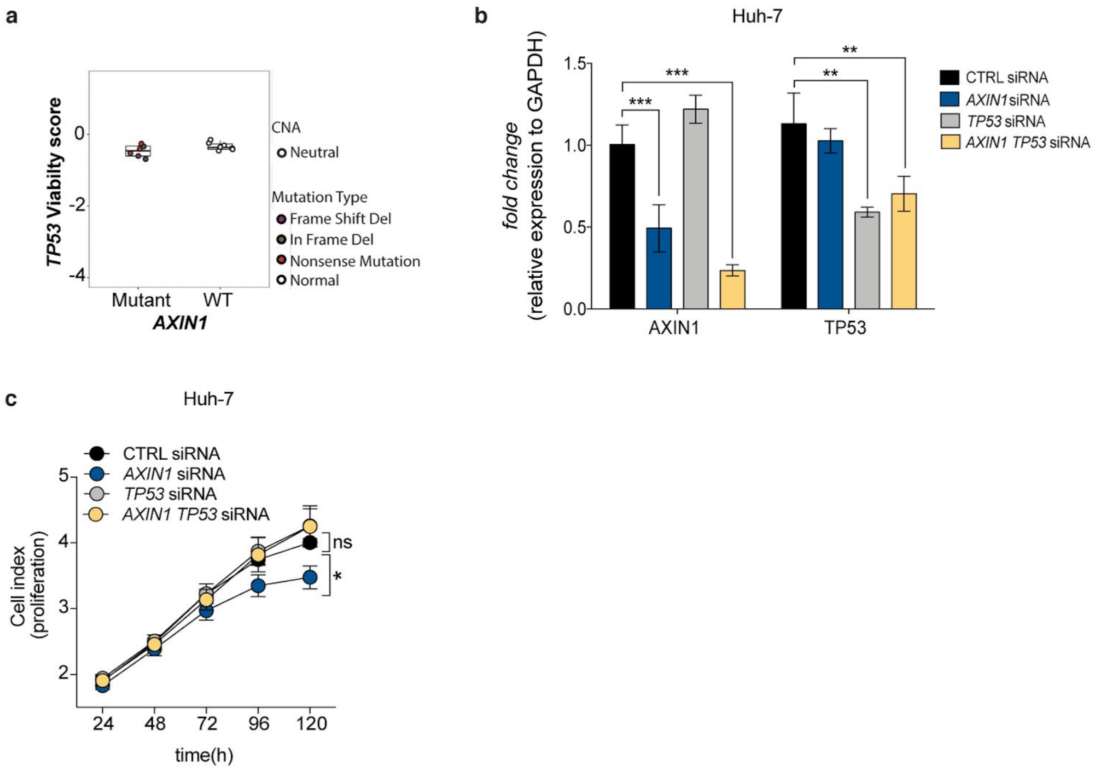


**Supplementary Fig. S2. Cross-validation of non-lethal interaction between *AXIN1* and *TP53* as negative control. a**, Viability scores of *AXIN1* mutant vs wild-type (WT) HCC cell lines with *TP53* knockdown. **b**, RNA expression levels (fold-change) of *AXIN1* and *TP53* relative to *GAPDH* in Huh-7 cell line transfected with control siRNA (black), *TP53* siRNA (gray), *AXIN1* siRNA (dark blue) or both (yellow). RNA levels were assessed by qPCR at 72 hours post siRNA transfection. Error bars represent SD from two independent experiments. **c**, Cell proliferation assay in Huh-7 cell line (*AXIN1* WT) transfected with control siRNA (black), *TP53* siRNA (gray),

*AXIN1* siRNA (dark blue) or both (yellow). Error bars represent SD from two independent experiments.

**Outlook for causal inference**

It should be noted that the average causal effect using matching is determined only from a subset of data. Since the mutation matrix is sparse, matching yields under-powered subsets in certain cases, inadequate for estimation of the average causal effect. We also tried to use IPTW (Inverse probability treatment weighting) method, but the results were unstable likely due to extreme weights resulting from the sparsity of the data and violation of the positivity assumption. Further, our current implementation is not conservative and is prone to selection bias, as we adjust for all the potentially confounding drivers without accounting for the underlying dependency structure between them. Therefore, it is important to learn the dependencies between the mutations and the outcome in the form of a directed acyclic graph (DAG). In the current implementation, we do not have this step due to small sample sizes. Based on these factors, we conclude that the consequent extension of our work would be to improve the estimation of the causal effect of driver mutations from sparse mutation data. We can also make use of knowledge from other omics data to select the confounders to adjust for, thus improving the overall estimate.
